# The crowding tax in parasites: linking dispersal behaviour to population growth in entomopathogenic nematodes

**DOI:** 10.64898/2026.10.25.761660

**Authors:** Ashwini Ramesh, Theresa Jones, Rayann Dorleans, Lexie Totaro, Farrah Bashey-Visser

## Abstract

1. Per capita fitness in group-living animals declines beyond an intermediate optimum, yet populations can overshoot this threshold if each newcomer still fares better in the group than alone. The resulting crowding tax (also known as Sibly tax) has been documented across free-living taxa, all of which retain the option of leaving a resource patch.
2. Parasites face a more constrained version of this trade-off. Once inside a host they cannot redistribute, and any overshoot in group size is fixed for the full reproductive bout. Whether macroparasites overshoot their fitness optimum, and what that costs the population, has remained difficult to quantify because it requires data on how parasites disperse among hosts, how they are distributed once there, and what individual fitness each of those burdens yields, all from the same system.
3. To tackle these challenges, we trace the causes and consequences of parasite dispersal behavior from individual-scale fitness to population-scale growth, using lab-experiments, mesocosms and field patterns paired with a model. We test these ideas in two species of entomopathogenic nematodes, *Steinernema costaricense* and *S. kraussei*, as a model system.
4. In mesocosms with standardized, equidistant hosts, both species dispersed as directional cohorts, generating heavy aggregation through this intrinsic behaviour. Dose-response experiments of infection, emergence and offspring production showed that per capita fitness peaks at low group sizes and drops sharply under the crowding that dispersal delivers, costing individuals 82% of their peak fitness in *S. costaricense* and 34% in *S. kraussei*.
5. Folding these fitness curves into an *R*_*0*_ model revealed that both species aggregate beyond the level that maximises population growth, realising 72% and 40% of their achievable *R*_*0*_. Applied to aggregation measured in a yearlong field survey, the model predicts that natural *S. kraussei* populations continue to pay this cost while natural *S. costaricense* populations, which aggregate far less, sit close to their own optimum.
6. Parasite aggregation in this system functions as a dispersal constraint that imposes a quantifiable fitness tax at the population level. Because the size of the tax depends on where the typical burden falls on each species’ offspring curve, two species with similar dispersal behaviour can pay very different prices. The tension between individual and group interests that is central to social foraging theory therefore extends to macroparasites.

## 1 INTRODUCTION

How organisms distribute among resource patches determines individual performance and population dynamics. Typically, group-living animals distribute themselves at intermediate densities that weigh the costs of crowding against the costs of isolation (Krause & Ruxton 2002). Theory predicts that *per capita* fitness should thus peak at intermediate densities. Yet the peak has different consequences for the individual and the population. Each “newcomer” benefits from joining even after the group has passed the optimum, because their fitness in a crowded group still exceeds fitness alone (Sibly 1983). This overshoot imposes a tax on every member of the group (also referred to as the Sibly tax; Sibly 1983), and has been quantified in foraging efficiency in baboons (Markham et al. 2015), nest-site competition in colonial birds (Brown 1969), and food intake rates in social foragers broadly (Giraldeau & Caraco 2000; Krause & Ruxton 2002). This tax, however, is not inevitable as most free-living animals can reassess and redistribute when crowding becomes too severe.

Parasites face the same trade-off but with a critical constraint: once they enter a host, they cannot redistribute. Here, the host is the patch and the infection intensity is the group size. Thus, the cost of overshooting is fixed for the entire reproductive cycle. Within hosts, this intensity governs parasite fecundity, body size, and survival (Poulin 2007). Because intensity varies among hosts, these individual-level costs are not shared equally. The most consistent distributional pattern in macroparasite ecology is aggregation, where few hosts contain most parasites and most hosts contain few or none (Crofton 1971; Shaw & Dobson 1995; Wilson et al. 2002). While aggregation is known to have profound consequences for host fitness and regulation of host populations (Hudson et al. 1998), less focus has been directed toward parasite populations themselves. If parasites overshoot the per capita optimum group size, as other group-living organisms do, it reduces not only individual fitness but population growth itself. Thus, how, and to what extent, does the cost of aggregation scale from individual parasite behavior to population dynamics?

Several challenges have limited our ability to connect parasite aggregation behavior to its fitness and population-level consequences. *First*, although parasites aggregate among hosts, overdispersion alone does not reveal the behavioral mechanism that produced it (Wilber et al. 2017). Clumping can arise passively from spatial heterogeneity and mass action (Keymer & Anderson 1979; Wojan et al. 2025), from individual parasites foraging optimally (Fretwell & Lucas 1969), or from group movement (Carlesso et al. 2024; Zhang & Lauder 2024). These three behavioral mechanisms can predict identical patterns when hosts differ in quality or position. Separating the parasite’s role requires hosts that are standardized, equidistant, and stationary, so that any departure from uniform spread arises from how parasites move. *Second*, aggregation among hosts places parasites across a range of within-host densities with opposing fitness consequences. At high densities, parasite per capita fecundity typically declines with parasite burden (Anderson & Schad 1985; Walker et al. 2009). At low densities, parasites typically fail to overcome host defences or find mates (May 1977). These opposing effects can produce an intermediate optimum where per capita fitness peaks. However, fitness components such as infection probability, body size, and fecundity respond to density with different nonlinear functional forms and can indicate different optima when measured separately. Integrating these components jointly is critical to understanding the fitness cost of the observed distribution. *Finally*, these within-host fitness costs scale to the population through the among-host distribution. Models of host-parasite dynamics have long incorporated aggregation as a parameter that modulates *R*_0_ (Anderson & May 1978; Wilber et al. 2021; Sun et al. 2026). Empirical tests, however, have focused on the host side, for instance, parasite-induced mortality in red grouse (Hudson et al. 1992) and parasite-reduced fecundity in Svalbard reindeer (Albon et al. 2002). In contrast, evaluating the cost from the parasite’s perspective requires knowing how many individuals experience each within-host density. Per capita fitness at each density, integrated over the among-host distribution, determines *R*_0_. Thus, evaluating whether and to what extent aggregation increases or reduces parasite *R*_0_ requires within-host density dependent fitness and among-host distribution data from the same system, integrated through a common model.

To tackle these challenges, we trace the causes and consequences of parasite dispersal behavior from individual-scale fitness to population-scale growth, using two species of entomopathogenic nematodes as a model system. These nematodes complete a full reproductive cycle within a single insect host in weeks, infection densities can be precisely controlled, hosts can be standardized by mass and species, and every offspring that emerges can be counted. Two species with differing dispersal abilities (*S. costaricense*, a highly mobile cruise forager; and *S. kraussei*, a less mobile forager) provide a contrast in how foraging behaviours may influence aggregation-fitness relationship. First, we quantify how these species behaviorally distribute among standardized, equidistant hosts in controlled soil arenas, testing whether the observed pattern is consistent with mass action, optimal foraging, or group movement. Second, we measure the fitness consequences of group size across the full range of doses, combining these into a net per capita fitness function. Finally, we integrate these fitness functions with the observed among-host distribution through a next-generation *R*_0_ calculation to ask whether observed aggregation maximizes, reduces, or has little effect on parasite population growth. We express *R*_0_ at the observed aggregation as a share of the maximum *R*_0_ attainable for each species, which tests directly whether aggregation is near optimal, costly or inconsequential. Thus, we are able to connect parasite dispersal behavior to individual-scale fitness to population-scale consequences in a single empirically parameterized framework.

## 2 MATERIALS AND METHODS

### Study System

To link parasite dispersal behaviour to its fitness consequences, we use two species of entomopathogenic nematodes (EPNs) in the genus *Steinernema*: *S. costaricense* and *S. kraussei* infecting larvae of the greater wax moth *Galleria mellonella*. EPNs are soil-dwelling insect parasites with a prolonged free-living juvenile stage (Strong et al. 1996). The free-living infective juveniles (IJs) are non-feeding and developmentally dormant but actively seek insect hosts (Bal and Grewal 2015). On entering a host, IJs resume development, kill the host within a week, and reproduce within the carcass. Offspring emerge as a new cohort of IJs a few weeks later and disperse into the soil (Selvan et al. 1993). This compact, single-host life cycle means that IJs emerge from a single cadaver as a point source, so the spatial distribution among hosts is determined entirely by nematode dispersal behaviours, though abiotic factors such as water movement and soil disturbance may also play a role. The fitness consequences of nematode behavior, including infection success, adult body size, and offspring production, can then be measured within a single generation in one host.

### 2.1 Part A: Parasite Behaviour — Dispersal and Distribution Among Hosts in Soil Mesocosm Arenas

To characterize aggregation behaviour during dispersal, we tested three mechanisms that predict distinct among-host distributions. (1) *Mass action:* parasites contact hosts in proportion to their relative encounter rates, so mean burden scales with dose per host number and variance equals the mean (Poisson). (2) *Optimal foraging:* parasites assess hosts and settle so as to equalise fitness across them, so burdens converge on a common value and variance falls far below the mean regardless of treatment (3) *Aggregated dispersal*: parasites travel as cohorts and colonize hosts together, so burdens are overdispersed with variance far exceeding the mean (negative binomial).

To discriminate among these hypotheses, we manipulated both parasite density and host availability, generating treatments with different mass-action predictions, and asked whether parasites track those predictions or distribute themselves according to the other two hypotheses. We released nematodes from a central point source, mimicking emergence from a single cadaver, into circular soil arenas (140 x 20 mm Petri dishes) containing a 50:50 mix of twice autoclaved local soil and MetroMix®820 (Sungro), maintained at 20% moisture. The design was a fully crossed 2 x 2 factorial with two levels of nematode density (1,000 or 10,000 IJs per arena) crossed with two levels of host availability (1 or 6 hosts per arena), with 12 replicate arenas run separately for each species. *G. mellonella* larvae (0.18 to 0.28 g) were each placed in a mesh pouch to limit movement, so that the spatial pattern of infection reflected parasite behaviour rather than host relocation. In 6-host arenas, hosts were arranged in a circle at equidistant positions 5 cm from the arena center and their position (1-6) recorded. Nematodes were pipetted to the center as a concentrated suspension in 1ml of dH20. Arenas were incubated at 20°C for 72 h, after which time all hosts were transferred from the soil to 60 x 20 mm petri dishes lined with filter paper, where hosts were checked for mortality daily for 7 days. Hosts were dissected at 96 h post-mortality. All nematodes were counted and classified as male, female, or immature, and body length of up to 10 females and 3 males per host was measured using an ocular micrometer on a dissecting scope. The final dataset comprised 318 dissected insect cadavers (*n* = 165 *S. costaricense, n* = 153 *S. kraussei*). Twelve fewer *S. kraussei* hosts were recovered for dissection. Uninfected (zero-nematode) hosts — 11 *S. costaricense* (6.7%) and 27 *S. kraussei* (17.6%) — were retained in all Part A analyses. Infection intensity and mean female size per host were modelled separately for each species using negative-binomial GLMs (MASS::glm.nb), with nematode density and host availability as fixed factors.

#### Quantifying Aggregation

To distinguish among the three dispersal mechanisms, we used the relationship between variance and mean in infection intensity. Variance equal to the mean indicates mass action (Poisson), variance below the mean suggests optimal foraging, and variance exceeding the mean indicates aggregation. We quantified this relationship at two scales. Across all hosts per species, we fit negative binomial distributions to infection counts pooled across treatments using maximum likelihood estimation (fitdistrplus package), extracting the aggregation parameter *k* and mean *μ*. A lower *k* indicates stronger aggregation. Within individual six-host arenas, we computed Ives’ (1991) *J* index (*J* = *V* /*μ*^2^ - 1/*μ*), where *V* = sample variance across the six hosts and *μ* the calculated mean across the 6 hosts. *J* = 0 corresponds to Poisson dispersion, *J* > 0 indicates aggregation, and *J* < 0 indicates uniformity. We estimated 95% confidence intervals on *J* using Tukey’s jack-knife (Ives 1991). For comparison, we also computed *J* from the single-host arenas, where among-arena variance reflects variation among independent point-source releases. If within-arena *J* exceeds among-arena *J*, then aggregation arises from parasite behaviour during dispersal and not merely from variation in infection success (the number of nematodes establishing per host) among independent releases that received the same total inoculum.

#### Spatial Orientation

Finally, aggregation may also have a spatial signature within arenas. Pheromone-mediated trail following (Erdogan et al. 2021) predicts that nematodes disperse directionally from a point source rather than radiating symmetrically. Directional dispersal should produce a strong spatial gradient in infection, whereas random dispersal should show no directional pattern. To distinguish these patterns, we reoriented infection counts within each six-host arena. Position 1 was assigned to the most-infected host and subsequent positions were numbered in the direction of the second-most-infected host, wrapping around the circle. This reorientation preserves the relative spatial orientation while removing the arbitrary starting position. To quantify directional dispersal, we also mapped reoriented positions onto angular distances from the most-infected host: position 1 = 0°, positions 2 and 6 = 60°, positions 3 and 5 = 120°, and position 4 = 180°. A negative relationship between infection (log(x + 1) transformed) and angular distance indicates a directional gradient. We assessed significance with a within-arena permutation test (1000 permutations), shuffling host positions within each arena and recomputing the regression slope to generate a null distribution. We report *R*^2^ and permutation *p*-values for each species–density combination (Fig. S2). Uninfected hosts (zero counts) were retained in all Part A analysis, including the infection-intensity GLM (Fig 1A), the pooled distribution (Fig 1B), the *J*-index (Fig 2A), and the log(x+1) spatial analysis (Fig 2B).

**Figure 1.**
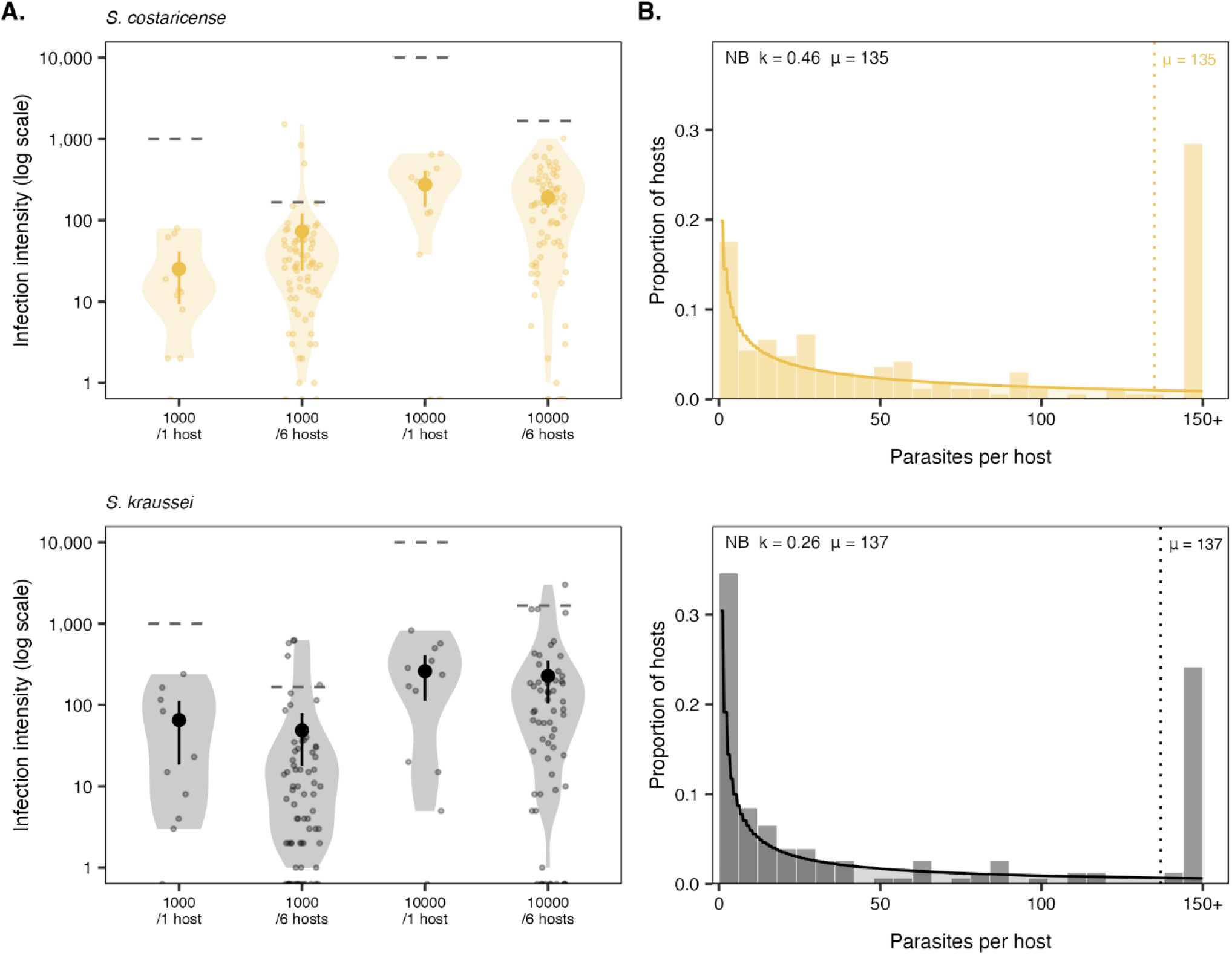
Distribution of parasites among hosts is strongly aggregated under all experimental combination of host and parasite densities. A. Treatment-level mean infection intensity (total nematodes per host) for *S. costaricense* (gold, top) and *S. kraussei* (black, bottom). Violin plots show the distribution of individual host values; filled circles with error bars are observed means with 95% CIs. Dashed horizontal lines show the relative mass-action predictions (nematode dose / number of hosts). B. Pooled negative binomial (NB) distributions across all treatments for each species. Histograms show the proportion of hosts in each burden class on a linear x-axis (counts exceeding 150 are pooled into the rightmost bin); curves are fitted NB density functions estimated by maximum likelihood. Dotted vertical lines mark the observed mean burden (*μ*). Inset text shows NB parameters (*k, μ*). Both species show strongly right-skewed distributions consistent with aggregation well beyond Poisson expectation.

**Figure 2.**
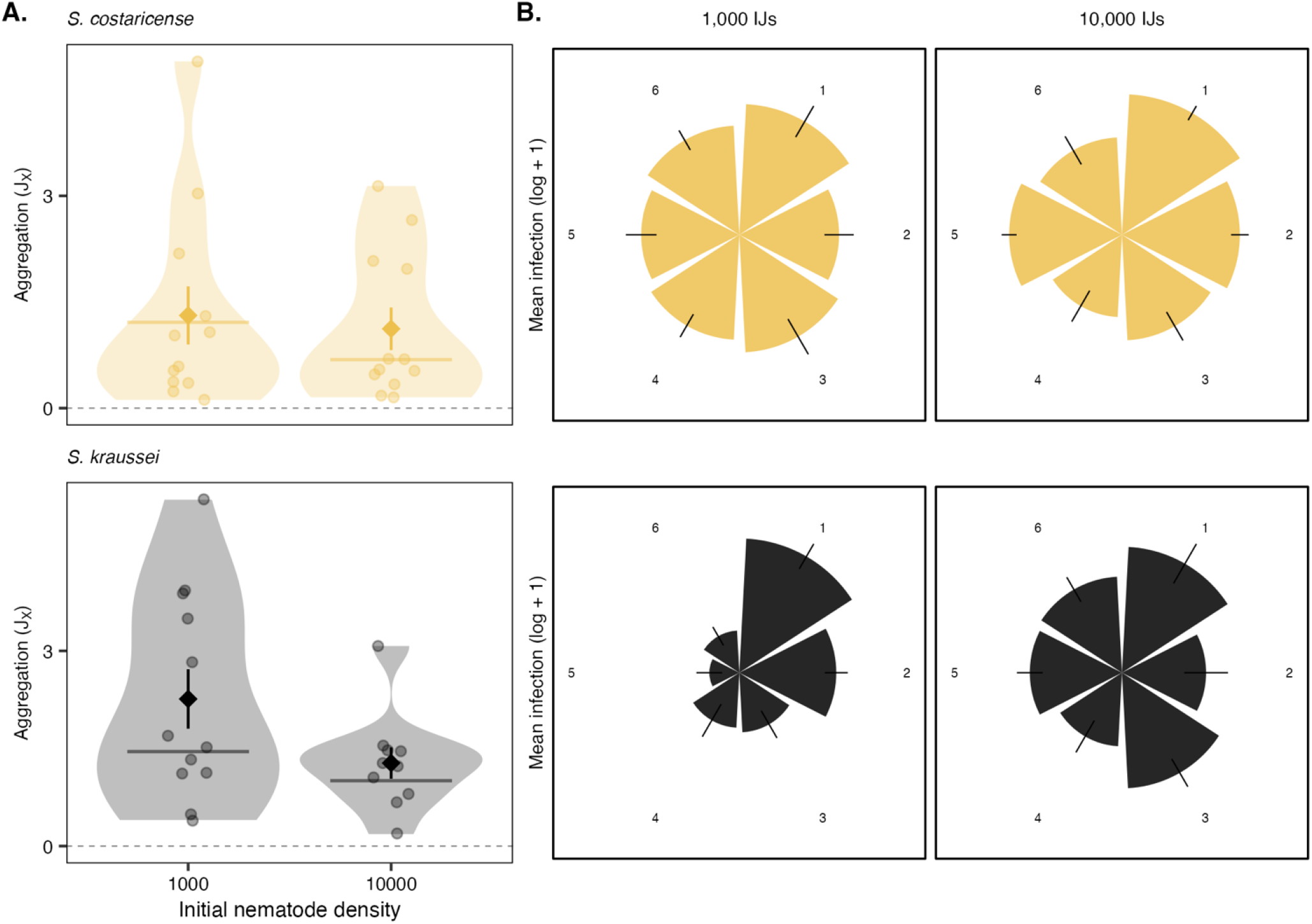
Within-arena aggregation exceeds among-arena variation, reflecting clumped nematode infections across hosts within arenas. A. Intraspecific aggregation index (*J*) for six-host arenas at two nematode densities, where each point is one arena. Diamonds with error bars are means with standard errors. Horizontal lines show the *J* index calculated across single-host arenas at the same density (the among-arena baseline). *J* > 0 indicates aggregation beyond Poisson expectation. Both species show strong positive *J* at both densities, with *S. kraussei* more aggregated than *S. costaricense*. Within-arena *J* exceeded the among-arena baseline in all four species-by-density combinations, iindicating that aggregation arises at least partly during dispersal within arenas. B. Polar rosette plots showing mean infection intensity (log(x + 1) transformed) by reoriented host position for 1,000 IJs (left) and 10,000 IJs (right). Position 1 is the most-infected host. Subsequent positions proceed around the arena in the direction of the second-most-infected host. The declining gradient from position 1 outward indicates directional cohort dispersal from the inoculation point rather than symmetric diffusion. See Fig. S2 for angular distance quantification with separate positional means.

### 2.2 Part B: Parasite Fitness—Infective-Dose – Reproductive-Output Assays on Single Hosts

Dispersal behaviour determines how many parasites each host receives, and this infection intensity shapes parasite fitness. To quantify the fitness consequences across a range within-host densities, we conducted a controlled dose-response experiment. Individual *G. mellonella* larvae were exposed to one of five IJ doses (20, 50, 100, 250, or 500 nematodes per host), plus unexposed controls for each species across three experimental blocks with 20 hosts exposed per dose per species in each block. Half the hosts were transferred to white traps upon death (Kaya and Stock 1997; Bashey & Lively 2009) to collect emerging offspring. The remaining hosts were dissected 72 hours post infection to determine infection status, count all nematodes and measure length for up to five adult females. Two datasets follow from this experiment, and because dissection requires destructive sampling they share no hosts and are linked only through the applied dose. (i) The dissected hosts were used to model establishment, establishment counts, and female body size (*S. costaricense, n* = 78; *S. kraussei, n* = 72). (ii) The white-trap hosts were used to model emergence, offspring number, and offspring size (*S. costaricense, n* = 81; *S. kraussei, n* = 64).

We modelled two fitness components, emergence probability *p*(*D*) and total offspring production *F*(*D*), each as a continuous function of dose, selecting the best fit by AIC. Emergence probability, the proportion of white-trap hosts that yielded any offspring, followed a complementary log-log relationship with dose (*D*), *p*(*D*) = 1 - exp(-exp(*a* + *b* * *D*)). The two parameters were estimated by maximum likelihood from the binary emergence outcome of each white-trap host. Emergence rather than establishment is the gate carried into the *R*_0_ model, because the pathway to the next generation requires the exit of transmission stage propagules from the host and not merely entry into it. (2) Total offspring production was modeled for hosts that produced offspring with zero-anchored points at dose = 0 to capture the origin. The relationship followed a Ricker function: *F*(*D*) = *a* · *D* · exp(−*b* · *D*), the mechanistic form predicted by scramble competition (Ives 1988). The Ricker can capture a hump-shaped relationship (peaking at dose 1/*b*) between dose and offspring number. Note, hosts that failed to produce offspring were not included in this dataset as they were handled by the emergence function (1 above). For both curves, 95% confidence intervals were estimated from 500 bootstrap resamples. Four additional fitness components, establishment probability, establishment counts, female body size, and offspring body size, are reported in the Supplementary Material (Table S3, Fig. S3).

### 2.3 Part C: Population-Level Consequences — Aggregation and R0

At the population scale, the reproductive output of a parasite depends on the full distribution of burdens across hosts. *R*_0_, the expected number of offspring per parasite per generation, was therefore obtained by integrating the single-host fitness functions from Part B over the negative binomial distribution of burdens (Churcher et al. 2005, 2006), rather than by evaluating them at the mean burden (Anderson and May 1978, 1985). The emergence probability and Ricker offspring function replaced the assumed fecundity functions of earlier models. These empirical functions already include any mate limitation over the doses tested, which is expected mainly at lower doses (Bashey and Lively 2009; Fig. S5B), so no separate mating term was added (May 1977). Because each generation is completed within a single host, *R*_0_ was computed by direct summation over burdens.

We modeled one complete parasite generation akin to our mesocosm experiment where hosts were made synchronously available. IJs emerge from a cadaver, disperse, infect new hosts in groups of size *G*, and produce offspring. The number of parasites each host receives is drawn from a negative binomial distribution with parameters *k* (aggregation) and *μ* (mean burden), estimated from Part A. For each group size *G*, the expected contribution to the next generation is *p*(*G*) * *F*(*G*), the probability that the host yields offspring times the total offspring produced. Averaging this contribution across the full distribution of group sizes yields the population-level reproductive number

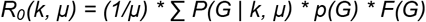

where *P*(*G* | *k, μ*) is the probability of a group of size *G*. Here *p*(*G*) is the emergence probability and *F*(*G*) the Ricker offspring function, both estimated from the white-trap hosts, so that *p*(*G*) · *F*(*G*) is the expected offspring production of an exposed host. *R*_0_ was evaluated across a grid of *k* from 0.05 to 1000 and summarised as efficiency, the value of *R*_0_ at the observed *k* as a proportion of the maximum *R*_0_ attainable over that grid; efficiency is bounded between 0 and 1 and does not depend on an interior optimum existing. Confidence intervals were obtained from 400 bootstrap iterations resampling the arena burdens, the emergence outcomes, and the offspring counts independently. The negative binomial is fitted to established burdens whereas *p* and *F* were measured against applied dose; the sensitivity of the result to this correspondence is examined in Fig. S5A.

At the individual level, per-capita fitness *w*(*G*) = *p*(*G*) * *F*(*G*) / *G* peaks at a group size *G*\* that represents the individually optimal infection intensity. For G*, p(G) is the establishment probability fitted with a curve anchored at zero, p(G) = 1 − exp(−a · G). If the observed mean burden *μ* exceeds *G*\*, the Sibly tax, or the individual’s cost for the population’s aggregated distribution, can be calculated as the fraction of maximum per-capita fitness lost to crowding (Sibly 1983). At the population level, we computed *R*_0_ across a grid of *k* values at the observed mean burden for each species. Because *R*_0_ approaches a plateau rather than an internal peak in one species, the *k* that maximises *R*_0_ is not reliably estimable (Fig. S4B). We therefore summarised the result as efficiency, the value of *R*_0_ at the observed *k* as a proportion of the maximum *R*_0_ attainable across the grid. Efficiency measures how much of the population growth a species realises under the distribution its dispersal actually produces.

#### Field Validation

To test whether laboratory predictions hold under natural conditions, we used field data from a year-long soil survey at Indiana University Research and Teaching Preserve at Moores Creek, Indiana (Ramesh et al 2024). Six 4 x 4 m forest sites were arrayed along an elevational gradient and sampled monthly by qPCR (see Ramesh et al 2024, for full protocol). For each site-month we estimated the negative binomial *k* by the method of moments, *k* = *μ*^2^ / (*V* − *μ*), where *μ* and *V* are the mean and variance of nematode counts, retaining only site-months where the sample variance exceeded the mean (*V* > *μ*), since *k* is undefined otherwise. Of the 72 site-month combinations, this yielded 64 estimates for *S. costaricense* and 60 for *S. kraussei*. Each field *k* was then evaluated on that species’ own *R*_0_ curve and expressed as efficiency, placing field and mesocosm populations on the same scale.

#### Software

All analyses were conducted in R (R Core Team 2024), using fitdistrplus for negative binomial fitting (Delignette-Muller and Dutang 2015), MASS for generalized linear models (Venables and Ripley 2002), pscl for zero-inflated models (Zeileis et al. 2008), emmeans and car for planned contrasts and Type III tests, and ggplot2 for figures (Wickham 2016).

## 3 RESULTS

### 3.1 Part A: Parasite Behaviour — Dispersal and Distribution Among Hosts in Soil Mesocosm Arenas

Nematodes distributed among hosts in a strongly aggregated pattern under all experimental conditions. As nematode density increased, both species showed a significant increase in the number of nematodes that infected each insect host (Fig 1A, Table S1). For *S. costaricense* in single-host arenas, the mean infection intensity at 10,000 vs. 1,000 IJs was approximately 11-fold higher, roughly matching mass-action scaling, while for *S. kraussei* infection intensity was 4-fold higher (Fig 1A, Table S1). Concomitant with this increase in infection intensity, female size decreased with nematode density for both species — females were approximately 1.2 mm smaller at 10,000 than at 1,000 IJs in *S. costaricense* and approximately 0.7 mm smaller in *S. kraussei* (Fig S1, Table S2). In contrast, the species differed in how they responded to host availability. For *S. costaricense*, at low nematode density adding hosts increased per-host burden, contrary to expectation based on mass action (Fig 1A, Table S1c,d). While at high nematode density adding hosts had little effect on per-host burden, such that there was a significant interaction between nematode density and host availability (LR *χ*^2^=5.29, *df* =1, *p* = 0.021, Table S1b). In contrast for S. kraussei, neither host availability nor its interaction with nematode density significantly affected infection intensity (Table S1b).

Under mass action, per-host infection should scale proportionally with nematode density per host, such that the infection intensities for the 6-host treatment should be lower than that 1-host treatment for each density. As described above, this prediction was not upheld: the only significant effect of host density was in the opposite direction. Additionally, the mass action prediction of random dispersal was not seen in the variance-to-mean ratio. While the Poisson expectation would be unity, the observed variance-to-mean ratio ranged from 31 to over 1000 across treatments (Table S1d). High variance-to-mean ratios reflect both aggregation among infected hosts and the substantial fraction of uninfected hosts (up to 21% in the 6-host *S. kraussei* treatment). These high variances also indicate that infection intensities were inconsistent with optimal foraging, which predicts equal infection burdens across hosts and near-zero variance. Instead, both species showed right-skewed distributions consistent with aggregated dispersal: many hosts lightly infected, few hosts heavily burdened (Fig 1B). When examining hosts pooled across all treatments, *S. kraussei* was more strongly aggregated than *S. costaricense* (*k* = 0.26 vs. 0.46, where lower *k* indicates stronger aggregation) despite similar mean burdens (*μ* = 137 vs. 135; Fig. 1B).

The intraspecific aggregation (*J*) index confirmed that aggregation arises during dispersal within individual arenas (fig. 2A). *J* exceeded zero in every species-by-density combination (jackknife 95% CIs excluded zero for all four groups) and was consistently higher for *S. kraussei* than *S. costaricense*, matching the stronger pooled aggregation (*k* = 0.26 vs. 0.46). Within-arena *J* also exceeded the among-arena baseline computed from single-host arenas (Fig. 2A). Aggregation therefore arises at least in part from the dispersal process within arenas, in addition to variation in infection success among independent releases that received the same total inoculum. To formally compare aggregation between species, we tested whether *J* indices differed at each density level and bootstrapped the difference in pooled *k. J* indices did not differ significantly between species at either density (Welch t-test: *p* = 0.136 at 1,000 IJs, *p* = 0.693 at 10,000 IJs), reflecting high arena-to-arena variability seen in Figure 2A. However, a bootstrap comparison of the pooled negative binomial *k* (1,000 iterations) confirmed that the species differ at the distributional level: *S. kraussei* had consistently lower *k* (more aggregated) than *S. costaricense* (difference = −0.207, 95% CI: −0.313 to −0.113). The species thus differ in their population-level aggregation pattern even though individual arenas are noisy. Because inoculum dose was held constant across arenas, this arena-to-arena variability — and the stronger aggregation of *S. kraussei* — is likely to reflect difference in the species’ dispersal behavior and their ability to overcome host defences.

Aggregation can be visualized in the six-host arenas by plotting the average number of nematodes that infected each host arranged spatially relative to the most-infected host (Fig. 2B). The polar rosettes show the clear dominance of the most-infected host (position 1), consistent with directional cohort dispersal rather than symmetric diffusion from the center point. Quantifying this pattern as angular distance from the peak host (Fig. S2), infection declined significantly with angular distance for three of the four species–density combinations: *S. kraussei* at 1,000 IJs (*r* =-0.45, *p* < .001), *S. costaricense* at 10,000 IJs (*r* =-0.32, *p* = .004), and *S. kraussei* at 10,000 IJs (*r* =-.198, *p* = .022). Only *S. costaricense* at 1,000 IJs showed no significant gradient (*r* =-0.13, *p* = .120). *S. kraussei* at low dose showed the strongest spatial structure, where nearly the entire cohort travelled in one direction. At higher nematode densities, the spatial gradient weakened but remained significant, consistent with multiple subgroups radiating in different directions while still generating the overall overdispersion that the *J* index detects (Fig. 2A). The species difference mirrors the negative binomial result as *S. kraussei* showed stronger spatial gradients than *S. costaricense* (e.g., *r* =-0.45 vs. -0.13 at 1,000 IJs), matching its stronger aggregation (lower *k*). Overdispersion among hosts thus arises from directional cohort dispersal.

### 3.2 Part B: Parasite Fitness—Infective-Dose – Reproductive-Output Assays on Single Hosts

The arena experiment demonstrated that both species produce strongly aggregated distributions among hosts, with infection intensities varying widely depending on a host’s position relative to the main dispersing cohort. To quantify the fitness consequences of this density variation, we conducted controlled dose-response experiments on single hosts across a range of doses. The probability that an exposed host yielded any offspring declined with dose in both species, and steeply in *S. costaricense* (Fig. 3A); the decline in *S. kraussei* was shallower and noisier. Establishment fell less steeply than emergence (Fig. S3A). At 500 IJs, half of the dissected *S. costaricense* hosts contained established nematodes whereas a fifth of white-trap hosts yielded offspring, so crowding acts mainly on reproduction.

**Figure 3.**
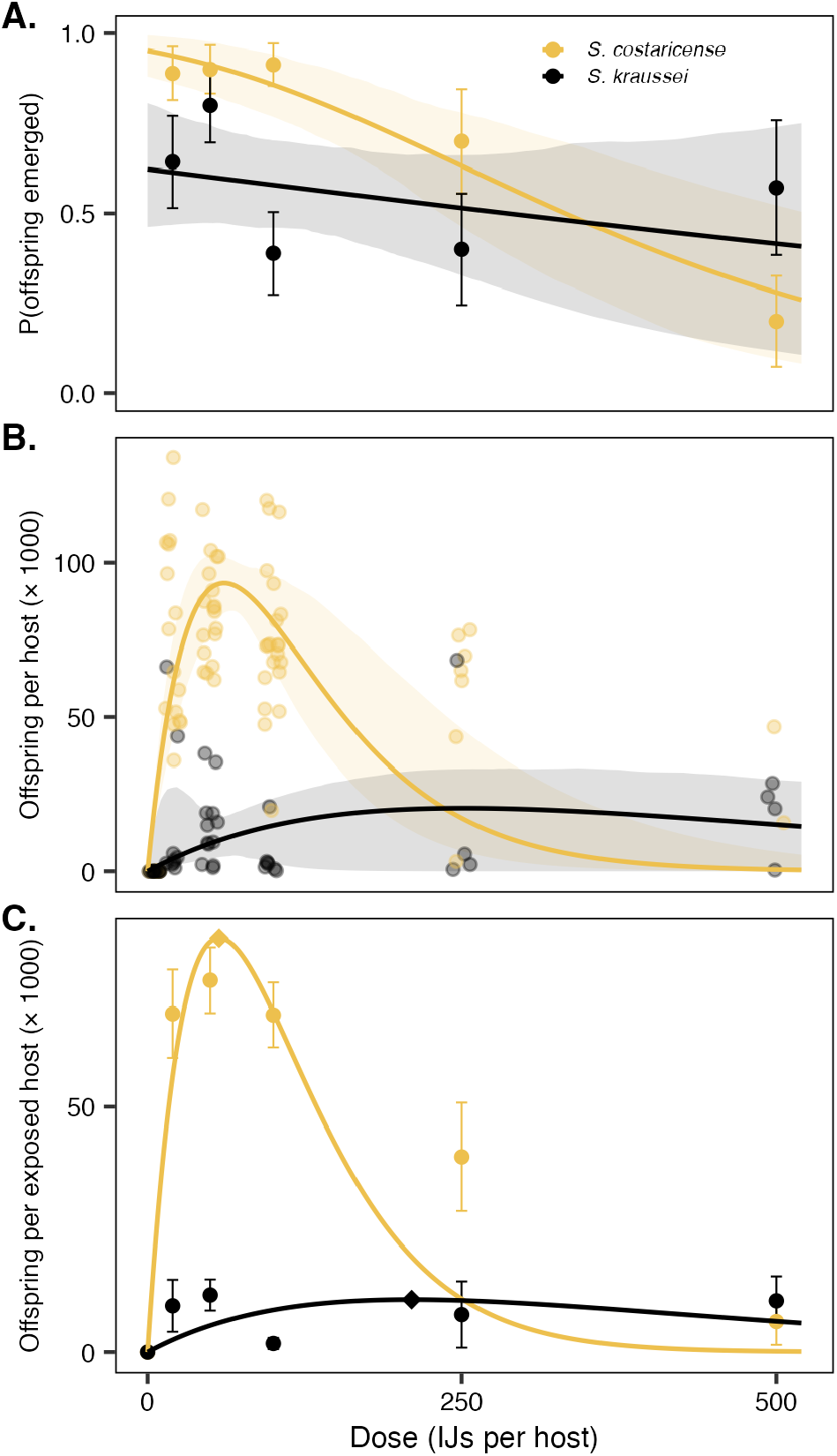
Crowding within a host reduces parasite fitness in two ways. A. Probability that an exposed host yielded emerging nematode offspring, scored on white-trap hosts. This declined with dose in both species, and steeply in *S. costaricense*, so that heavily exposed hosts frequently produced no transmission stage parasites. B. Total offspring per host among hosts that successfully produced emerging nematodes, fitted with a Ricker function. *S. costaricense* (gold) and *S. kraussei* (black) are shown in all panels. Points are calculated proportions ± 1 binomial standard error (A) or the number of parasites emerging from individual hosts (B); shading shows 95% bootstrap confidence intervals. C. Expected offspring per exposed host, the product of the curves in A and B and the quantity the *R*_0_ model sums. Points are means ± 1 standard error over all white-trap hosts, zeros included, and diamonds mark the peaks at 57 IJs (*S. costaricense*) and 210 IJs (*S. kraussei*). Fitted parameters are given in Table S3. Establishment probability, establishment counts, female body size and offspring body size are shown in Fig. S3.

Total offspring produced per host followed a Ricker curve in each species (Fig. 3B, Table S3). *S. costaricense* peaked sharply at a dose of 61 IJs and dropped off dramatically thereafter, indicating strong negative density dependence on offspring production. *S. kraussei* showed a more gradual peak with density, increasing in offspring production up to a dose of approximately 250 and showing only a minor decline at higher densities. Additionally, offspring size declined slightly with dose for *S. kraussei* (*R*^2^ = 0.124, *p* = .038) but showed no trend for *S. costaricense* (*R*^2^=0.028, *p* = .18; Fig. S3D). Expected offspring per exposed host, which combines emergence and offspring number and is the quantity the *R*_0_ model sums, peaked at 57 IJs in *S. costaricense* and 210 IJs in *S. kraussei* (Fig. 3C). In sum, the cost of crowding within a host is manifest as lower parasite reproductive success.

### 3.3 Part C: Parasite Population Growth — Aggregation and R0

We used the empirically fitted functions from Part B to derive a single per-capita fitness function, *w*(*G*), for each species to explore the effect of group size (*G*) on per-capita fitness and population growth. *w*(*G*) = *p*(*G*) · *F*(*G*) / *G*, where *p*(*G*) is the probability that a host receiving *G* parasites becomes infected, fitted with a curve anchored at zero, and *F*(*G*) is the number of offspring a productive host yields. Per-capita fitness peaked at group sizes well below the observed mean burden (Fig. 4A). *G*\* = 21 IJs for *S. costaricense* and *G*\* = 26 IJs for *S. kraussei* (Fig. 4A); bootstrap resampling confirmed the stability of the *S. costaricense* estimate, whereas the *S. kraussei* estimate was less certain because its crowding parameter is weakly constrained (Fig. S4A). The observed mean burden *μ* exceeded *G*\* by a factor of 6 for *S. costaricense* and a factor of 5 for *S. kraussei* (Fig. 4A, dotted vertical line). The Sibly tax, the fraction of maximum per-capita fitness lost to crowding, was 82% for *S. costaricense* and 34% for *S. kraussei* (Fig. 4A, vertical arrow).

Given that the distribution concentrates parasites far from the individual optimum, we asked if this mismatch also reduces population-level fitness (*R*_0_)? Integrating per-capita fitness over the negative binomial distribution yields *R*_0_ as a function of the aggregation parameter *k* (Fig. 4B). *R*_0_ rose steeply as aggregation weakened. For *S. costaricense* it reached a shallow maximum at *k* = 4.0 before flattening, whereas for *S. kraussei R*_0_ increased monotonically towards an even distribution with no interior optimum within the range examined. Both observed distributions of k correspond to lower *R*_0_ values. *S. costaricense*, at *k* = 0.46, achieves 72% of the maximum its own fitness curves allow (95% CI 54–87%); *S. kraussei*, at *k* = 0.26, achieves 40% (95% CI 34–80%). Both species therefore pay a population-level cost for aggregating, and *S. kraussei*, which aggregates more intensely, pays the steeper one.

**Figure 4.**
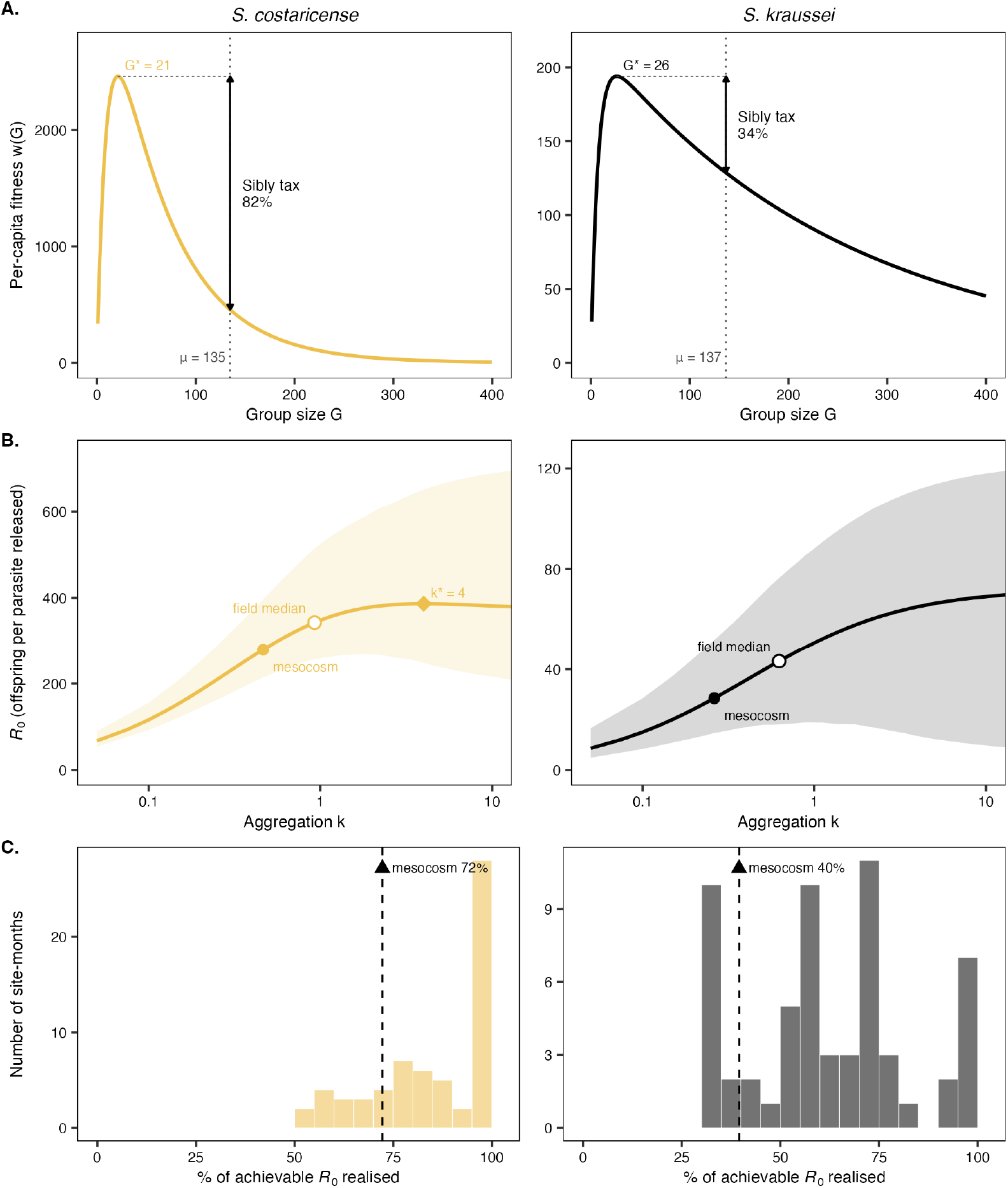
Aggregation costs population growth, and the cost differs between species. Columns are *S. costaricense* (left) and *S. kraussei* (right). A. Per-capita fitness, *w*(*G*) = *p*(*G*) · *F*(*G*) / *G*, the expected offspring of one parasite in a host shared by G parasites, where p(G) is the probability of infection. The dotted vertical line marks the observed mean burden *μ*; the horizontal dashed segment is a guide carrying peak fitness across to *μ*; the vertical arrow is the Sibly tax, the fraction of peak per-capita fitness lost to crowding (82% and 34%). B. Population-level *R*_*0*_*(k, μ) = (1/μ) Σ P(G* | *k, μ) · p(G) · F(G)*, the expected offspring per parasite averaged over all hosts when burdens follow a negative binomial with aggregation k and mean μ, where p(G) is the probability that a host yields offspring, shown as a function of k at fixed mean burden, on a logarithmic *k* axis, with the 95% bootstrap envelope shaded. *R*_0_ was evaluated over the full grid to *k* = 1000; the axis is truncated at *k* = 10, beyond which both curves are flat. Filled circles mark the mesocosm distribution; open circles the median field distribution and the diamond the *R*_0_-maximising *k* for *S. costaricense* (*k*\* = 4.0); *S. kraussei* has no interior optimum. Note the five-fold difference in absolute *R*_0_ between species. C. Distribution of the percentage of achievable *R*_0_ realised by each field site-month, obtained by evaluating the curve in B at that site-month’s estimated *k*; bars count site-months in each 5% bin (*n* = 64 and 60 site-months). The dashed line and triangle mark the mesocosm value.

#### R_0_ optimum

Whether aggregation can raise *R*_0_ depends on where the mean burden falls on the offspring curve (Fig. 3B). *S. costaricense* produced the most offspring at 61 IJs per host and declined steeply thereafter, consistent with rapid exhaustion of a single cadaver, whereas *S. kraussei* tolerated much higher densities and peaked at 252 IJs. Mean burdens were nearly identical in the two species (135 and 137 nematodes per host), so a typical *S. costaricense* host was already overcrowded whereas a typical *S. kraussei* host held fewer parasites than the peak. Aggregation therefore had opposite effects. In *S. costaricense*, uneven spread left some hosts with fewer parasites and closer to the peak, so *R*_0_ was highest at moderate aggregation (k = 4.0). In *S. kraussei*, uneven spread left most hosts with even fewer parasites, so *R*_0_ rose steadily as parasites became more evenly spread. More generally, uneven spread raises mean output only where the mean burden lies beyond twice the peak dose, 122 IJs for *S. costaricense* and 503 IJs for *S. kraussei*, because beyond that point the offspring curve bends upward. This comparison treats the burden counted in arena hosts and the dose applied in the dose-response experiment as the same scale. Only a fraction of applied nematodes established (Fig. S3B), so an arena host carrying a given burden was more crowded than a dose-response host given the same number, and Fig. S5A shows how efficiency changes across the range of mean burdens.

#### Field validation

Field populations were markedly less aggregated than mesocosm populations (median *k* = 0.93 for *S. costaricense, n* = 64 site-months; 0.62 for *S. kraussei, n* = 60 site-months), likely because heterogeneous soil conditions and variable time from nematode emergence to sampling reduce overdispersion. Evaluating each site-month *k* on its own species’ *R*_0_ curve, and expressing the result as the share of achievable *R*_0_ realised, separates the two species (Fig. 4C). *S. kraussei* populations, which were observed to be more aggregated, remained costly in the field, realising a median 60% of achievable *R*_0_ (interquartile range 51–75%), with only 7 of 60 site-months at or above 95%. *S. costaricense* populations, by contrast, realised a median 89% (interquartile range 75–98%), with 28 of 64 site-months at or above 95%. The model therefore predicts a population-level cost of aggregation in the field for *S. kraussei*, whereas many *S. costaricense* populations aggregate little enough to sit close to their own optimum.

## 4 DISCUSSION

How does parasite dispersal behavior shape the distribution of parasites among hosts, and what are the consequences of that distribution for individual fitness and population growth? Using soil mesocosm arena experiments and single-host dose-response assays, we traced the path from dispersal behavior to population growth in two entomopathogenic nematode species. Both species dispersed as directional cohorts from a central release point, producing strongly aggregated distributions inconsistent with mass action and optimal foraging predictions. This aggregation arose likely from intrinsic dispersal behavior as confirmed by spatial analyses showing significant directional gradients in infection intensity for three of four species–density combinations, consistent with pheromone-mediated group movement documented in other EPN systems (Shapiro-Ilan et al. 2014). Controlled dose-response experiments then revealed that offspring production per host peaks at an intermediate dose, 61 IJs for *S. costaricense* and 252 IJs for *S. kraussei*. Per capita fitness shows an even lower optimal infection size, suggesting an individual level cost of 34-82%. Integrating these fitness functions over the observed distributions showed that both species aggregate far more than the level that maximizes population growth, sacrificing 28% (*S. costaricense*) and 60% (*S. kraussei*) of their potential *R*_0_. This fitness tax is analogous to the group-size costs predicted by Sibly (1983) for free-living organisms, where individuals rationally joining a group pushes group sizes beyond the per-capita optimum. Applied to field aggregation from a year-long soil survey, the model predicts that natural *S. kraussei* populations pay this cost, whereas natural *S. costaricense* populations aggregate little enough to sit close to their own optimum (Ramesh et al 2024). Our study traces the complete path from individual dispersal behavior through individual fitness to population-level growth, demonstrating that parasite aggregation behaviour imposes a quantifiable fitness tax on their population growth.

Aggregation in both species arose from intrinsic dispersal behavior, independent of host heterogeneity. In nature, infective juveniles emerge *en masse* from a single cadaver as a concentrated pulse of offspring with no prior information about host location. With hosts standardized and uniformly positioned, both species still produced strongly overdispersed distributions that rejected both random dispersal and the ideal free distribution (optimal foraging), which predicts equal burdens across equivalent hosts. Observed variance far exceeded the mean in all treatments. Aggregation is near-universal in host-parasite systems (Shaw and Dobson 1995), but in most vertebrate-helminth systems the source is host heterogeneity in exposure or immune competence (Calabrese et al. 2011; Luong et al. 2011). Here we show that aggregation persists even when host heterogeneity is removed, consistent with intrinsic aggregation documented in EPN species dispersing through homogeneous arenas without host cues (Shapiro-Ilan et al. 2014; Kaplan et al. 2020; Stevens et al. 2024). The angular pattern of infection revealed how this aggregation forms (Fig. 2B, Fig S2). Infection declined significantly with angular distance from the most-infected host in three of four treatments, consistent with a cohort of parasites moving directionally like a wave through the arena rather than a diffusing cloud radiating symmetrically from the release point. The host in the path of this wave received the heaviest infection and those on the far side of the arena received nearly nothing. The two species differed in the tightness of this wave. *S. kraussei*, which has more limited dispersal abilities, a longer lifespan (Bashey et al. 2016) and strong host chemotaxis (Enriquez et al, unpublished data), maintained stronger cohort cohesion during movement. Nearly the entire cohort travelled in one direction at low dose, producing the strongest spatial gradient and the strongest overdispersion (Figure 2). *S. costaricense*, a highly mobile cruise forager which shows no host chemotaxis, spread more broadly across hosts, showing weaker spatial gradients and less extreme aggregation. The two species also differed in their sensitivity to host availability. For *S. costaricense*, nematode density and host availability interacted significantly (Table S1b). Adding hosts at low-nematode density increased per-host burden, while at high-nematode density more hosts slightly reduced per-host burden. This difference is consistent with cohorts of nematodes dispersing at random with respect to hosts. At low nematodes densities, more cohorts (and hence more individuals) are successful at finding a host as more hosts are available. For *S. kraussei*, host availability did not affect infection intensity, as this species can sense and move towards hosts, but were limited in their ability to reach hosts before the mesocosm trial ended. Thus, both species were found to show an aggregated distribution among host, with the degree of this aggregation consistent with intrinsic differences in mobility and host sensing.

The fitness consequences of an aggregated distribution depend on the effects of within-host density on parasite survival and reproduction. Controlled dose-response experiments revealed a nonlinear relationship between infective dose and total per-host offspring production (Fig 3B). At high doses, competition for finite cadaver resources likely depresses reproduction as nematodes share a fixed resource pool (Selvan et al. 1993; Koppenhoefer and Kaya 1996). At low doses, total output rises because more parasites share the host, even though offspring per parasite was highest at the lowest dose tested. Facilitation may still occur at very low doses, where infections can fail to establish the symbiotic bacteria (Goodrich-Blair & Clarke 2007) or to include both sexes (Bashey & Lively 2009). Density-dependent fecundity of this kind is general across helminth systems (Michael and Bundy 1989; Walker et al. 2009; Neves et al. 2021), but in vertebrate helminths the mechanism can be immune-mediated (Paterson & Viney 2002). In EPNs the immune response is suppressed early in infection (Castillo et al. 2011), so resource competition within the cadaver likely drives the density dependence documented here. The two species experience this tradeoff differently. *S. costaricense* peaked sharply at low dose, producing far more offspring per host at its optimum but crashing steeply under crowding. *S. kraussei* produced fewer offspring at its optimum but maintained reproductive output across a fourfold wider range of densities. This broader tolerance of crowding is consistent with *S. kraussei* being the more aggregated species, since a species that routinely experiences high within-host densities would be expected to tolerate crowding better. Despite these contrasting strategies, per-capita fitness peaked at similar group sizes for both species (Figure 4A), far below the mean burden observed in arenas. Integrating per capita fitness over the observed negative binomial distribution shows that neither species is distributed in the way that would maximise its own population growth. *S. costaricense* realises 72% of its achievable *R*_0_ and *S. kraussei* 40%. The size of the shortfall is set jointly by the aggregated distributions that dispersal creates and by the hump-shaped relationship between dose and offspring production, so the same behaviour carries a different price in the two species.

This overshoot is consistent with Sibly’s (1983) prediction that group sizes exceed the per-capita optimum, a principle well-documented across group-living organisms from social foraging birds to baboon troops (Beauchamp 1998; Giraldeau and Caraco 2000; Markham et al. 2015). A mechanistically closer parallel comes from carrion fly communities, where blowflies arrive at carcasses in pulses and compete for a finite resource that cannot be redistributed. Aggregation of flies on carcasses reduces per-capita fitness, and variance in group size across carcasses governs competitive outcomes through Jensen’s inequality (Ives 1991). Here we find that the same principle applies to parasites. Earlier models established that *R*_0_ depends on aggregation (Anderson and May 1978; Sun et al. 2026) and density-dependent fecundity (Churcher et al. 2005), but both relied on assumed functional forms rather than measured fitness components. An infective juvenile that joins a crowded host still gains higher infection probability than one that disperses alone, and the population bears the cost. Unlike flies and other social foragers, however, parasites cannot leave a host once they have entered it, so this cost cannot be corrected after infection. How large the cost becomes depends on the shape of each species’ fitness curve. *S. costaricense*, with its steep fitness curve, pays primarily through fecundity competition in crowded hosts, whereas *S. kraussei*, with its flatter curve, pays through parasites spread across hosts too lightly infected to reach the dose at which offspring production peaks. Whether aggregation can help at all is set by where the mean burden falls on the offspring curve, and for *S. costaricense* it falls just past the point at which variance begins to pay. The cost arises because dispersal overshoots even this more permissive optimum. Field populations from the year-long soil survey suggest that both the cost and this species difference persist outside the laboratory. Natural populations were less aggregated than mesocosm populations, likely because heterogeneous soil and time since emergence from a cadaver partially reduce overdispersion. Yet field *S. kraussei* aggregation corresponds to only 60% of achievable *R*_0_, indicating that these populations aggregate beyond the optimum and likely pay the same fitness cost, whereas field S. costaricense populations, being far less aggregated, largely escape it. Thus, dispersal behavior is predicted to constrain population growth in both controlled and natural settings, and because the size of that constraint depends on how within-host density shapes each species’ reproductive potential, two similar species sharing a habitat can pay different costs of aggregation.

Several extensions could sharpen these conclusions. *First*, the experimental design standardized hosts to isolate parasite behavior. In nature, hosts vary in mass, immune status, and spatial distribution (Calabrese et al. 2011; Luong et al. 2011). Host heterogeneity could amplify aggregation by concentrating parasites on susceptible individuals or dampen it if parasites preferentially colonize unoccupied hosts. Adding host variation to the *R*_0_ model would test whether the fitness cost persists under more realistic conditions. *Second*, the cost of aggregation depends on how many parasites a typical host receives, and that number varies widely in nature. Across a wide range of mean burdens (Fig. S5A), *S. kraussei* falls short of its achievable *R*_0_ throughout, because its typical host stays below the dose at which offspring production peaks, whereas *S. costaricense* approaches its own optimum once burdens grow heavy enough for variation among hosts to place parasites near that peak rather than beyond it. Quantifying burdens in naturally infected hosts would show where each species sits on this curve. Because dissection is destructive, offspring production cannot be measured against the established burden of the same host, so our model treats arena burden and applied dose as one scale. *Third*, the shape of the fitness curve depends on the primary drivers of density dependence. Resource competition drives negative density dependence in EPNs, whereas immune limitation drives it in vertebrate helminths (Paterson and Viney 2002). Repeating the dose-response experiment in hosts that differ in size or immune competence would test whether a different mechanism shifts the *R*_0_ optimum. *Finally*, the *R*_0_ calculation presented here is a single-generation measure. Over multiple generations, aggregation can stabilize host-parasite dynamics (Anderson and May 1978), and the mechanism generating aggregation determines the degree of stabilization (Adler and Kretzschmar 1992; Rosà and Pugliese 2002). Multi-generation models that combine the Ricker fitness function with negative binomial dispersal would test whether a within-generation cost is offset by long-term persistence.

The tension between individually rational behavior and group-level outcomes is central to behavioral ecology (Sibly 1983; Giraldeau and Caraco 2000; Krause and Ruxton 2002), and the results presented here show that this tension extends to macroparasites. Two nematode species sharing a habitat (Ramesh et al 2024) pay very different prices for similar behaviours, because the cost of aggregation depends not only on the degree of aggregation but also on how per-host burden affects parasite fitness. Infective juveniles emerge from cadavers, follow pheromone trails, and infect hosts with conspecifics. Joining a group protects against infections too small to produce both sexes (Bashey and Lively 2009), but the resulting burdens reduce per capita fitness and population-level *R*_0_. Understanding whether the population-level cost of aggregation is a general feature of macroparasites will require connecting dispersal mechanisms to population dynamics across a wider range of host-parasite systems.

## AUTHOR CONTRIBUTIONS

AR (lead) and FB conceptualized the study and acquired funding. AR, TJ, RD, and LT carried out the experiment. AR analysed the data and wrote the first draft of the manuscript with major revisions from FB. All authors edited and approved the final draft of the manuscript.

## ACKNOWLEDGEMENTS

We thank Michael Wade’s “Population Genetics of Structured Populations” graduate class that inspired AR in framing key ideas for the paper. We thank E Huenupi and K Beidler for sharing valuable knowledge about soil sampling; M Chitwood for maintaining the IURTP properties. We are grateful to L Whitney, RK Phillips, SN Ramesh and S Ramesh for their assistance in the field sampling. AR thanks S Dubner, B Riley (‘Current’) and the Amalienburg for artistic inspiration.

